# Multiple P450s and Variation in Neuronal Genes Underpins the Response to the Insecticide Imidacloprid in a Population of *Drosophila melanogaster*

**DOI:** 10.1101/132373

**Authors:** Shane Denecke, Roberto Fusetto, Felipe Martelli, Alex Giang, Paul Battlay, Alexandre Fournier-Level, Richard A. O’Hair, Philip Batterham

## Abstract

Insecticide resistance is an economically important example of evolution in response to intense selection pressure. Here, the genetics of resistance to the neonicotinoid insecticide imidacloprid is explored using the Drosophila Genetic Reference Panel, a collection of inbred *Drosophila melanogaster* genotypes derived from a single population in North Carolina. Imidacloprid resistance varied substantially among genotypes, and more resistant genotypes tended to show increased capacity to metabolize and excrete imidacloprid. Variation in resistance level was then associated with genomic and transcriptomic variation, implicating several candidate genes involved in central nervous system function and the cytochrome P450s *Cyp6g1* and *Cyp6g2.* CRISPR-Cas9 mediated removal of *Cyp6g1* suggested that it contributed to imidacloprid resistance only in backgrounds where it was already highly expressed. *Cyp6g2*, previously implicated in juvenile hormone synthesis via expression in the ring gland, was shown to be expressed in metabolically relevant tissues of resistant genotypes. *Cyp6g2* overexpression was shown to both metabolize imidacloprid and confer resistance. These data collectively suggest that imidacloprid resistance is influenced by a variety of previously known and unknown genetic factors.

## Introduction

The introduction of synthetic insecticides is often followed by the appearance of resistance phenotypes in field populations, leading to significant reductions in agricultural production^1^. There has been much debate about whether the evolution of resistance is caused by variation in a single gene (monogenic) or by the additive effects of many (polygenic)^2,3^. Substantially more work has been dedicated to characterizing the monogenic variants, but such alleles arise in a genetic background where there is polygenic variation for tolerance to the insecticide^2^. Much still remains unclear about the relative contribution of different alleles to insecticide resistance, but *D. melanogaster* is uniquely placed to answer such questions, owing to the extensive genetic toolkit that has been developed in this model insect.

Imidacloprid is amongst the most widely used insecticides. It is derived from nicotine and is a member of the neonicotinoid chemical class. Neonicotinoids target nicotinic acetylcholine receptors (nAChRs) that have vital roles in neurotransmission and behaviour in insects^4,5^. However, imidacloprid resistance via mutations in targets is not the most common resistance mechanisms, possibly due to associated fitness costs^6^. The overexpression of cytochrome P450 enzymes (P450s) more frequently underpins imidacloprid resistance^7^. Some members of the P450 superfamily can function as drug metabolizing enzymes (DMEs) with xenobiotic substrates, while others have vital roles in development using endogenous substrates^8^. P450s which are capable of metabolizing imidacloprid and conferring resistance have been identified in several species^9–11^; the *Cyp6g1* gene of *D. melanogaster* has been particularly well studied^12^. Originally identified by mapping DDT resistance in the Hikone-R strain to a region containing a cluster of three P450 genes (*Cyp6g1*, *Cyp6g2* and *Cyp6t3*), resistance was shown to be due to the overexpression of *Cyp6g1*^13^. This overexpression was subsequently found to be caused by the insertion of the long terminal repeat of the retrotransposon, *Accord*, upstream of the gene^14^.

The expression of *Cyp6g1* is highly variable in field populations due to the *Accord* insertion, copy number variation and further transposable element insertions^15–17^. The ancestral M haplotype contains a single copy of *Cyp6g1* and expresses low levels of the gene compared to the more derived AA haplotype, which contains a duplication of *Accord-Cyp6g1* cassette in addition to several partial chimeric repeats of *Cyp6g1-Cyp6g2* which are not characterized^16^. Further modifications of the AA haplotype resulted from the insertions of the transposable element *HMS-Beagle* and a P element upstream of *Cyp6g1,* creating the BA and BP haplotypes, respectively. These derived haplotypes have been associated with increased levels of *Cyp6g1* expression and resistance to insecticides such as DDT and azinphos-methyl^16,17^. *Cyp6g2* expression correlates with *Cyp6g1* expression in the DGRP, but the contribution of this gene to resistance has not been shown^17^.

The structural modification of imidacloprid in biological systems includes both nitroreduction and oxidation reactions. Metabolites from both pathways have been detected in plants, animals and insects, but soil bacteria produce predominantly the nitroreduction metabolites^18–20^. In the case of *D. melanogaster,* the presence of nitroreduction metabolites is thought to be mostly due to endosymbiotic bacteria (Fusetto et. al this issue). Insect P450s are thought to produce oxidative metabolites exclusively, and the metabolites formed by CYP6G1 have been the best characterized. Heterologous expression of *Cyp6g1* in the tobacco plant *Nicotiana tabacum* produced the metabolites IMI-5-OH, IMI-Ole and IMI-diol^21^. These results were replicated when driving the expression of *Cyp6g1* in *D. melanogaster*^*22*^. The potential for the other 87 *D. melanogaster* P450s to be involved in imidacloprid resistance has yet to be tested; *Cyp6g1* is so far the only P450 linked to imidacloprid resistance in this species. While it is possible that no other *D. melanogaster* P450 is capable of metabolizing imidacloprid, this appears unlikely given that many P450s are polyspecific, and multiple P450s are often upregulated in field resistant insects^23,24^. Furthermore, the transcriptional response to xenobiotics is often regulated by transcription factors, such as *Cap ‘n’ Collar* and *DHR96,* that regulate the expression of many P450s^25,26^.

A common method of describing the genetic basis of a trait is to associate genetic variation with phenotypic variation, attempting to identify the causative quantitative trait loci (QTLs) and transcripts. This approach assumes no *a priori* knowledge about the genes that influence a phenotype, and applying these techniques in model organisms with well characterized genetic resources further enhances detection power. The Drosophila Genetic Reference Panel (DGRP) exemplifies these capabilities. The DGRP is a collection of 201 fully sequenced inbred *Drosophila* stocks, which represents a snapshot of genetic diversity present in a single population from Raleigh, North Carolina, sampled in 2012^27,28^. Using the DGRP, a genome wide association study (GWAS) can be performed by testing the associations of the ∼2.5 million genetic variants (most commonly single nucleotide polymorphisms; SNPs) across the DGRP genomes with phenotypic data for any quantitative trait. Further, sequencing of the DGRP male and female transcriptomes allowed for similar association studies to be performed with transcript expression level in a transcriptome wide association study (TWAS)^29,30^. The DGRP has been used to understand the genetic basis of a wide variety of traits, including insecticide resistance^17^.

Here, the genetic basis of imidacloprid resistance in the DGRP is described. The Wiggle Index (WI) bioassay^31^ was used to estimate imidacloprid resistance by measuring acute imidacloprid response, and substantial variation was observed among DGRP genotypes. Quantification of imidacloprid and its metabolites via high-performance liquid chromatography coupled with mass spectrometry (HPLC-MS) showed differences between resistant and susceptible subsets of the DGRP, suggesting that differences in overall imidacloprid metabolism significantly contribute to the differences in resistance. Many QTLs and transcripts were associated with imidacloprid resistance, implicating several genes involved in Central Nervous System (CNS) development as well as the P450s *Cyp6g1* and *Cyp6g2*. The subsequent deletion of *Cyp6g1* from two laboratory strains showed no significant differences in imidacloprid resistance, while the same deletion from a resistant DGRP genotype significantly decreased resistance. These deletions allowed for the direct measurement of the contribution of different *Cyp6g1* haplotypes to imidacloprid resistance. *Cyp6g2* was also linked to imidacloprid resistance in the DGRP by observing increased expression of the gene in the metabolically relevant tissues (midgut and Malpighian tubules) in resistant genotypes. Transgenic overexpression confirmed its ability to metabolize and confer resistance to imidacloprid. These data suggest that genetic variation in CNS development and the expression levels of *Cyp6g1* and *Cyp6g2* contribute to imidacloprid resistance in field populations of *D. melanogaster.*

## Results

### Analysis of the Distribution of Imidacloprid Resistance

Measurement of imidacloprid resistance in the DGRP was accomplished using the WI^31^, which measures the acute (one hour) motility response of third instar larvae to insecticide exposure, at two doses (25 and 100ppm). Imidacloprid resistance was quantified by relative movement ratios (RMRs), which reflect the motility of larvae after one hour of insecticide exposure relative to the motility of the same larvae before exposure (An RMR of 1 reflects no imidacloprid response, while an RMR of 0 implies the strongest possible response). Substantial variation in mean RMRs was observed among the 171 DGRP genotypes tested (Fig. 1; Supplementary Table S1; Supplementary Table S2). The mean RMRs of each genotype in the population showed significant correlation between the two doses (Adjusted R^2^=.18; p-value <3.6x10^-9^, Supplementary Fig. S1), but not between the amount of initial motility and final RMR (Adjusted R^2^=.001, p-value=.06; Supplementary Fig. S2). This suggests that the imidacloprid response was independent of larval motility measured in the absence of imidacloprid. The 25ppm exposure produced a slightly left skewed distribution of RMRs and is discontinuous due to 3 extremely susceptible genotypes. The 100ppm dose produced a more even distribution of RMRs. Broad sense heritability estimated for each dose was estimated to be 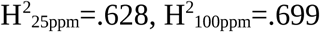.

**Figure 1-.**
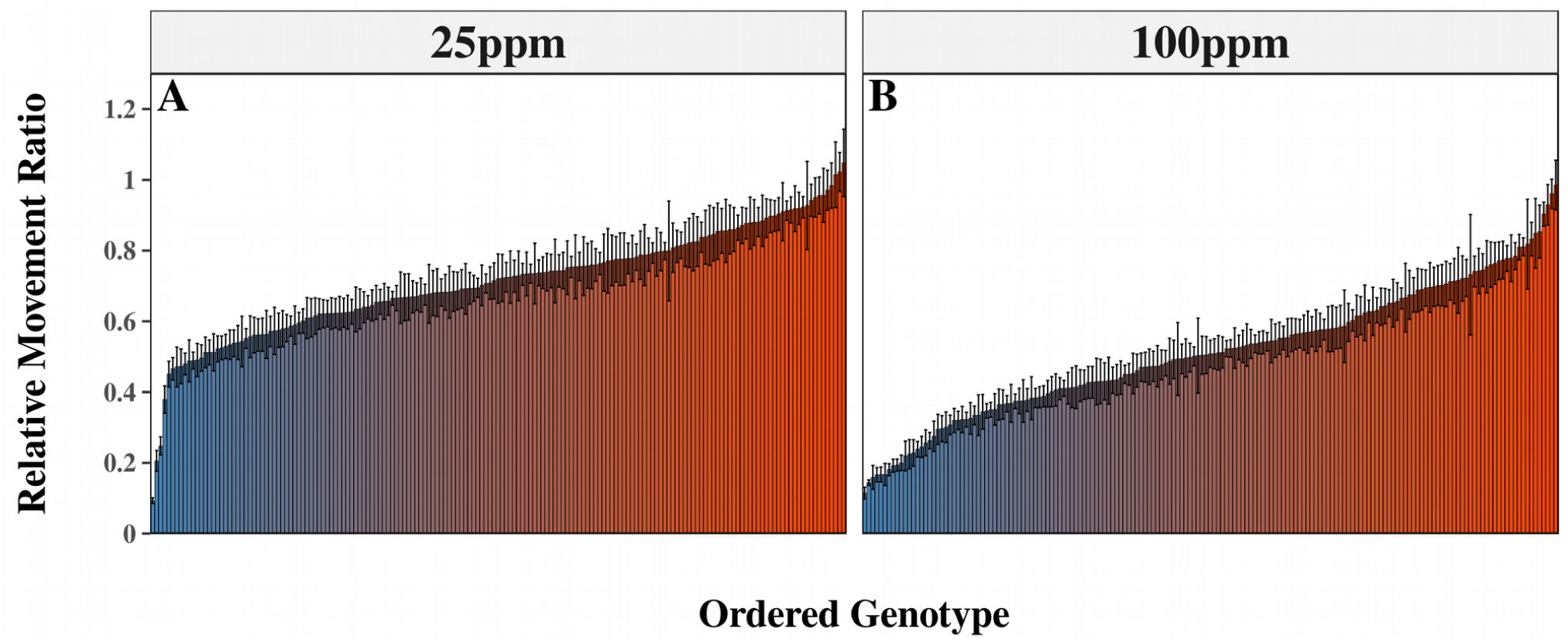
Phenotypic Spread of the DGRP. Imidacloprid response in the DGRP was assessed using the Wiggle Index at 2 doses A) 25ppm and 100ppm. Relative movement ratios represent the amount of imidacloprid response with a value of 1 reflecting no response and 0 the most substantial response. At each dose there was a spectrum of phenotypic responses ranging from susceptible (blue) to resistant (red). Error bars represent the standard error of the mean.

### Imidacloprid and Metabolite Quantification in the DGRP

To estimate the contribution of insecticide metabolism to the observed differences in imidacloprid response, the amount of imidacloprid and its metabolites (IMI-5-OH and IMI-Olefin) recovered from both larvae and the exposure media was quantified for resistant and susceptible subsets of the DGRP. This was performed under exposure conditions almost identical to those used to assess resistance in the DGRP (Fusetto et. al this issue), and metabolic phenotypes were tested for correlation with the imidacloprid response measured at 25ppm (Fig. 1A). The quantity of imidacloprid in larval bodies showed a significant positive correlation with RMR at 25ppm among genotypes. The more imidacloprid found in the body of a genotype, the stronger the imidacloprid response (Fig. 2A). Additionally, the quantities of IMI-5-OH and IMI-Olefin recovered from the media showed significant negative correlation with RMR, suggesting that increased levels of metabolites in the media provided for a weaker response to imidacloprid (Fig. 2 E,F). However, RMR did not significantly associate with the level of either metabolite in the body (Fig. 2 B,C). These data suggest that imidacloprid metabolism is higher in resistant genotypes and that these metabolites are preferentially excreted from the body.

**Figure 2-.**
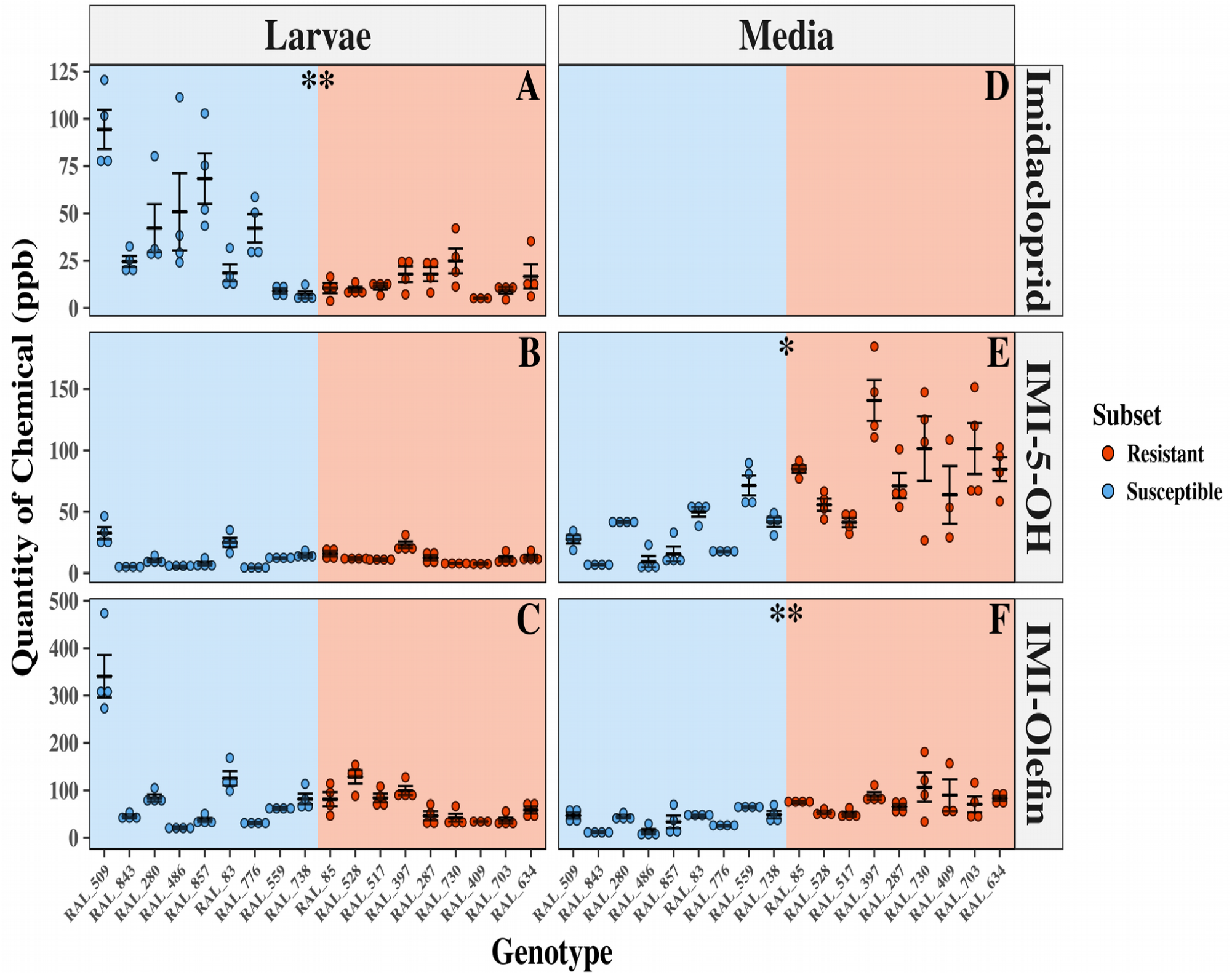
HPLC-MS in Resistant and Susceptible DGRP Subsets. The amount of Imidacloprid (A,D) IMI-5-OH (B,E) and IMI-Olefin (C,F) recovered from larval bodies (A-C) or the media (D-F) is reported in parts per billion (ppb) from a subset of the most susceptible (blue) and resistant (red) DGRP lines. No data is presented for imidacloprid in the media, due to the relative abundance of this molecule in the media, which makes detecting changes impossible. Error bars represent standard error of the mean. Stars represent the significance of association of each phenotype with RMR among genotypes using Pearson’s correlation test (*=p<.05; **=p<.001; ***=p<.0001).

### A GWAS for Imidacloprid Response Yields Many Neuronal Candidate Genes

A GWAS was performed in order to identify the genetic basis of imidacloprid resistance. The genome wide association of the scores (-log p-values) of annotated genetic variation in the DGRP was uniformly distributed, suggesting test-statistics were not inflated (Supplementary Fig. S3). Manhattan plots showed only 30 variants that crossed the P≤10^−5^ threshold and only one that fell below the Bonferroni threshold (Supplementary Fig. S4). Linkage disequilibrium between associated variants was low; only two minor linkage disequilibrium peaks were found among the associated variants.

The annotated function of the genes nearest to the significantly (P≤10^−5^) associated genomic variants implicated a high proportion of candidates having roles in the development and function of the CNS (Supplementary Table S3). 52.6% (10/19) of these genes are reported to have enriched expression in the third instar CNS (expressed 2 fold or greater compared to all other third instar tissues) compared to 19.6% when all *D. melanogaster* genes are considered. A number of the candidate genes have not been annotated, precluding Gene Ontology term analysis, but several genes appear to have described roles in CNS development or function. A single SNP near the S*ickie* gene was the only variant which elicited a p-value below the Bonferroni threshold (p=.04) at 25ppm and was also the only variant to be significantly associated at both 25ppm and 100ppm (Supplementary Table S3). These findings suggest that CNS development and function may contribute to imidacloprid response.

### A Transcriptome Wide Association Study Suggests Cyp6g1 and Cyp6g2 are Influence Imidacloprid Response

RNA-seq data for 185 DGRP genotypes^30^ was used to associate the expression level of specific genes with imidacloprid response. Unlike the GWAS candidate list, the TWAS candidate list was not enriched for genes expressed in any particular third instar tissue (Supplementary Table S4). Furthermore, no pattern emerged with regard to any process or function. However, the well known DME *Cyp6g1* was the most significant candidate at both doses (Fig. 3A, Supplementary Table S4). *Cyp6g2* expression was also significantly associated at 100ppm (Fig. 3B), and the expression of the two genes is highly correlated. There was no evidence that any variant from the GWAS was influencing the expression of any significantly associated transcripts, as no transcript eQTL was present among the significantly associated GWAS variants.

**Figure 3-.**
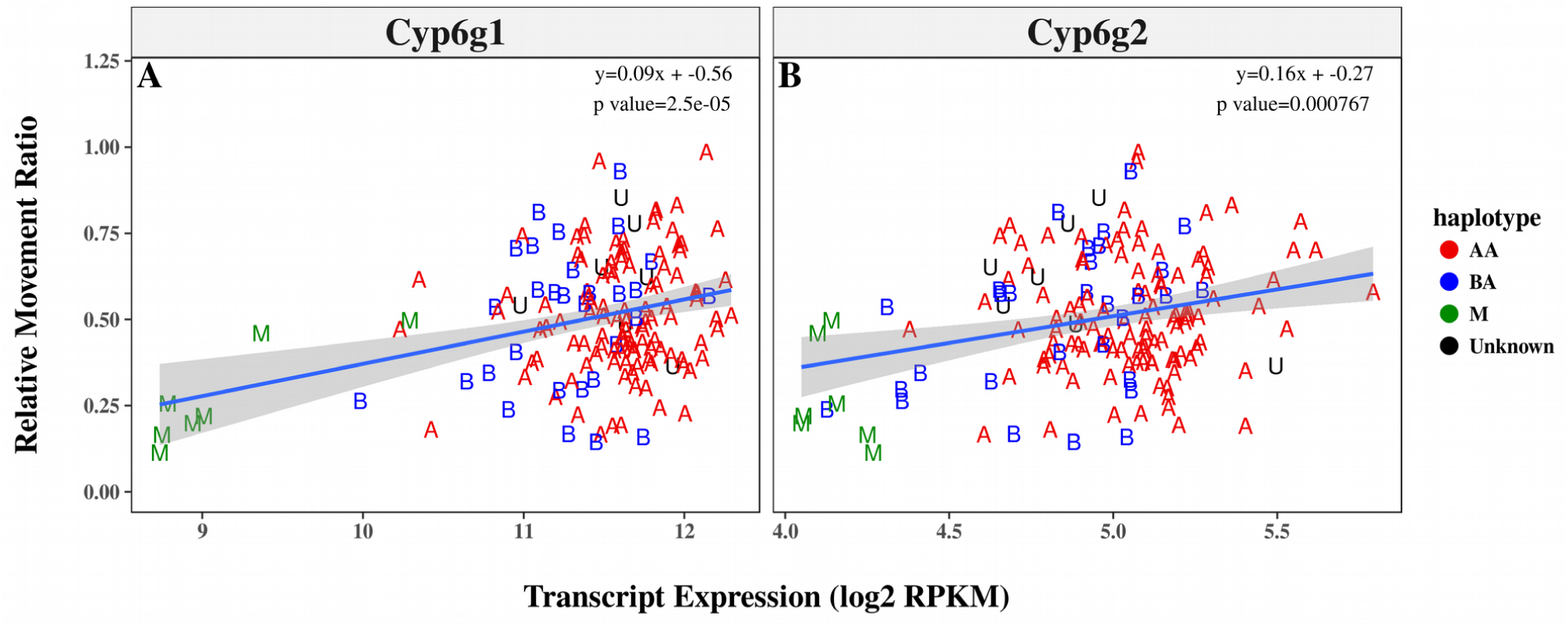
Transcriptional Association of Cyp6g1 and Cyp6g2. The association of the A) Cyp6g1 and B) Cyp6g2 transcripts with imidacloprid resistance is shown. Each plot compares the transcript’s expression reported by Huang et. al (2015) to the RMR of each genotype reported in the current study. Points are labelled according to their haplotype at the Cyp6g1 locus. A linear model is fit with a 95% confidence interval.

### The Knockout of Cyp6g1 Displays Haplotype Dependent Effects

The imidacloprid response of three *Cyp6g1* knockouts was compared to their matched controls in WI bioassay. RAL_517 showed significantly less imidacloprid response than RAL_517-Cyp6g1KO at both 25 and 100ppm (Fig. 4A,B). These findings were not replicated when testing knockouts in the Canton-S and Wxac backgrounds. No significant differences were found between these knockouts and controls when exposed to 5ppm imidacloprid, a dose used to detect potential response differences in the far more susceptible Canton-S and Wxac genotypes (Fig. 4 C-D). These data suggest that *Cyp6g1* makes a significant contribution to imidacloprid metabolism in BA haplotypes but not in backgrounds carrying an M haplotype.

**Figure 4-.**
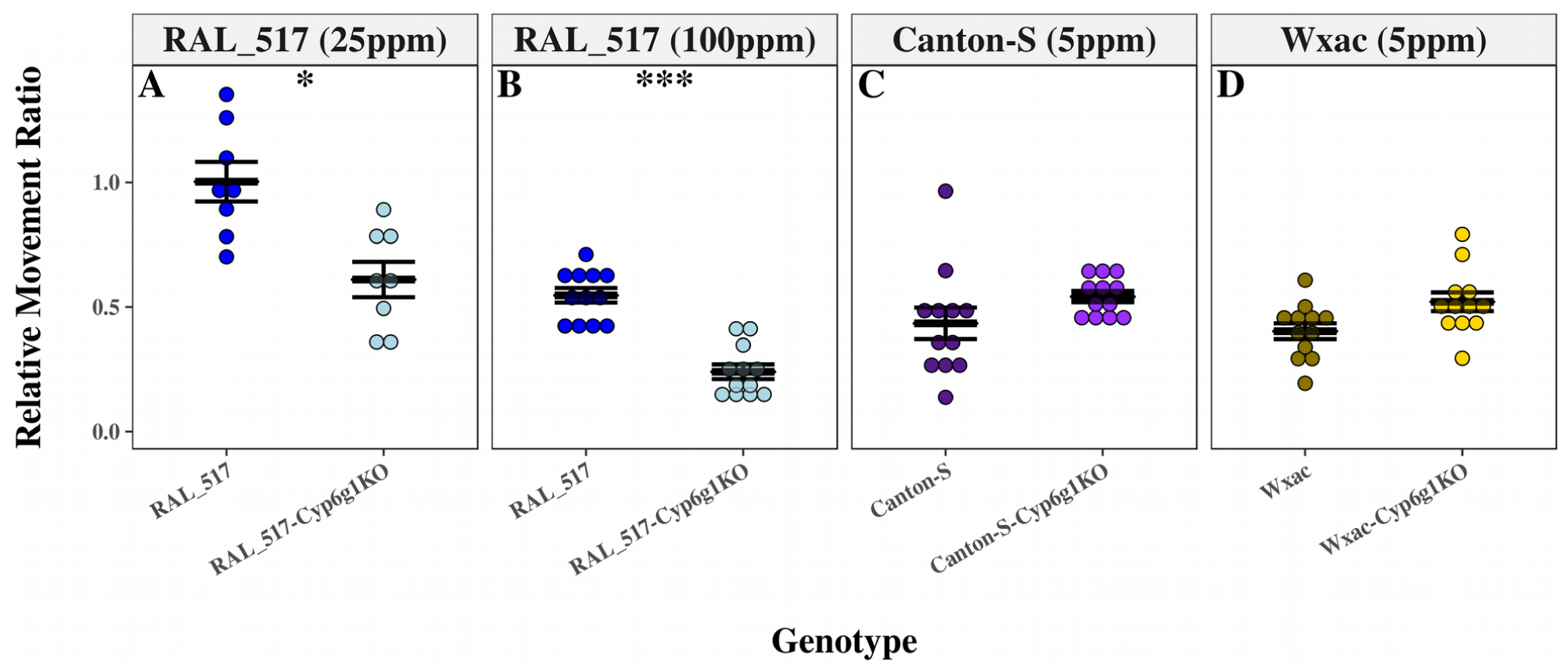
Imidacloprid Resistance in Cyp6g1KO Backgrounds. The effect of the removal of Cyp6g1 from 3 backgrounds on imidacloprid response is shown. Control lines are shown as dark colours while lighter colours represent knockouts. Removing Cyp6g1 from RAL_517 increased imidacloprid susceptibility at both A) 100ppm and B) 25ppm. No significant changes were observed when Cyp6g1 was removed from either Canton-S or Wxac using a discriminatory dose of 5ppm. Error bars represent the standard error of the mean. Stars represent the significance of the difference between the two genotypes measured by the Students T-test corrected for multiple comparisons with the Bonferroni method (*=p<.05; **=p<.001; ***=p<.0001).

### Cyp6g2 Expression is Enriched in the Digestive Tissues of Resistant Genotypes and Metabolizes Imidacloprid

The potential for the other P450 genes adjacent to *Cyp6g1* (*Cyp6g2* and *Cyp6t3*) to contribute to imidacloprid resistance in the DGRP was tested, by quantifying the expression of all three genes in the digestive tissues (midgut and Malpighian tubules) in a subset of DGRP genotypes (two AA and two M haplotypes). All samples showed consistent expression of the housekeeper gene *RP49* with the exception of the 3^rd^ and 4^th^ biological replicates of the RAL_360 genotype; these samples were excluded from the analysis. The remaining genotypes showed significantly higher levels of both *Cyp6g1* and *Cyp6g2* in the midguts and Malpighian tubules of AA haplotypes compared to M haplotypes (Fig. 5A-C; Supplementary Table S5). Although, the upregulation of *Cyp6g2* appears far weaker than for *Cyp6g1,* both genes showed larger differences between haplotypes when only the digestive tissues were considered compared to the whole body data reported previously (Fig. 5D-F, Supplementary Table S5)^30^. No significant patterns were observed for *Cyp6t3.* These data suggest that both *Cyp6g1* and *Cyp6g2* are upregulated in these digestive tissues in the AA haplotype, compared to the ancestral M haplotype.

**Figure 5-.**
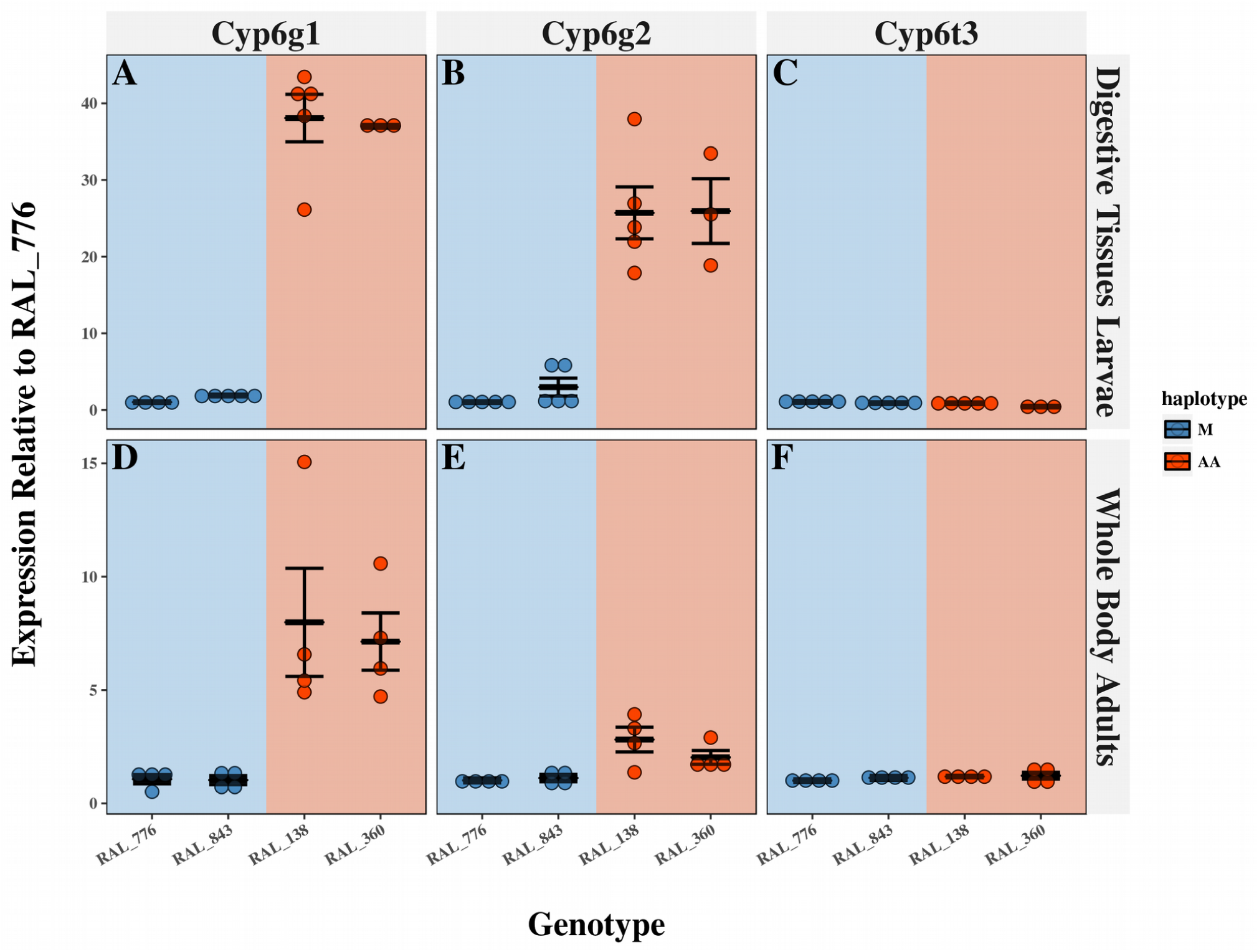
The Expression of P450s of the Cyp6g1 Locus in the Digestive Tissues. The expression of A) Cyp6g1 B) Cyp6g2 and C) Cyp6t3 was quantified in the digestive tissues of third instar larvae in 4 DGRP genotypes. M haplotypes are represented by blue points and AA haplotypes by red. The whole body expression data presented by Huang et. al (2015) is presented for the same genes D-F). Error bars reflect the standard error of the mean. Significant differences were detected between all Cyp6g1 and Cyp6g2 measurements between M and AA haplotypes (ANOVA Tukey’s honestly significant difference) with p-values reported in Supplementary Table S5.

The relative capacity of *Cyp6g1* and *Cyp6g2* to confer resistance to imidacloprid was tested by overexpressing the genes in the digestive tissues using the HR_GAL4 driver^14^ and two newly created UAS genotypes, which contained each gene’s open reading frame in a common insertion site on the second chromosome. Compared to their controls, genotypes overexpressing either *Cyp6g1* or *Cyp6g2* showed significantly higher resistance to imidacloprid, with the magnitude of resistance conferred by *Cyp6g1* being significantly higher (Fig. 6A,B; Supplementary Table S6). Although mRNA levels were not measured, the increased resistance conferred by *Cyp6g1* relative to *Cyp6g2* expressed from the same insertion site suggests that *Cyp6g1* enzyme may have a higher capacity to confer resistance to imidacloprid. To further verify the capacity of *Cyp6g2* to metabolize imidacloprid, HPLC-MS was employed to measure levels of imidacloprid and metabolites in a previously reported *Cyp6g2* overexpression genotype^32^. While the HR_GAL4 x w^1118^ control produced relatively low levels of metabolites and had high levels of imidacloprid in the body, HR_GAL4 x UAS-*Cyp6g2-3d* larvae produced higher levels of imidacloprid metabolites and had less imidacloprid in the body (Supplementary Fig. S5 A,E,F). These data indicate that *Cyp6g2* can act as a DME against imidacloprid, although it is possible that its capacity to confer resistance may be less than that of *Cyp6g1.*

**Figure 6-.**
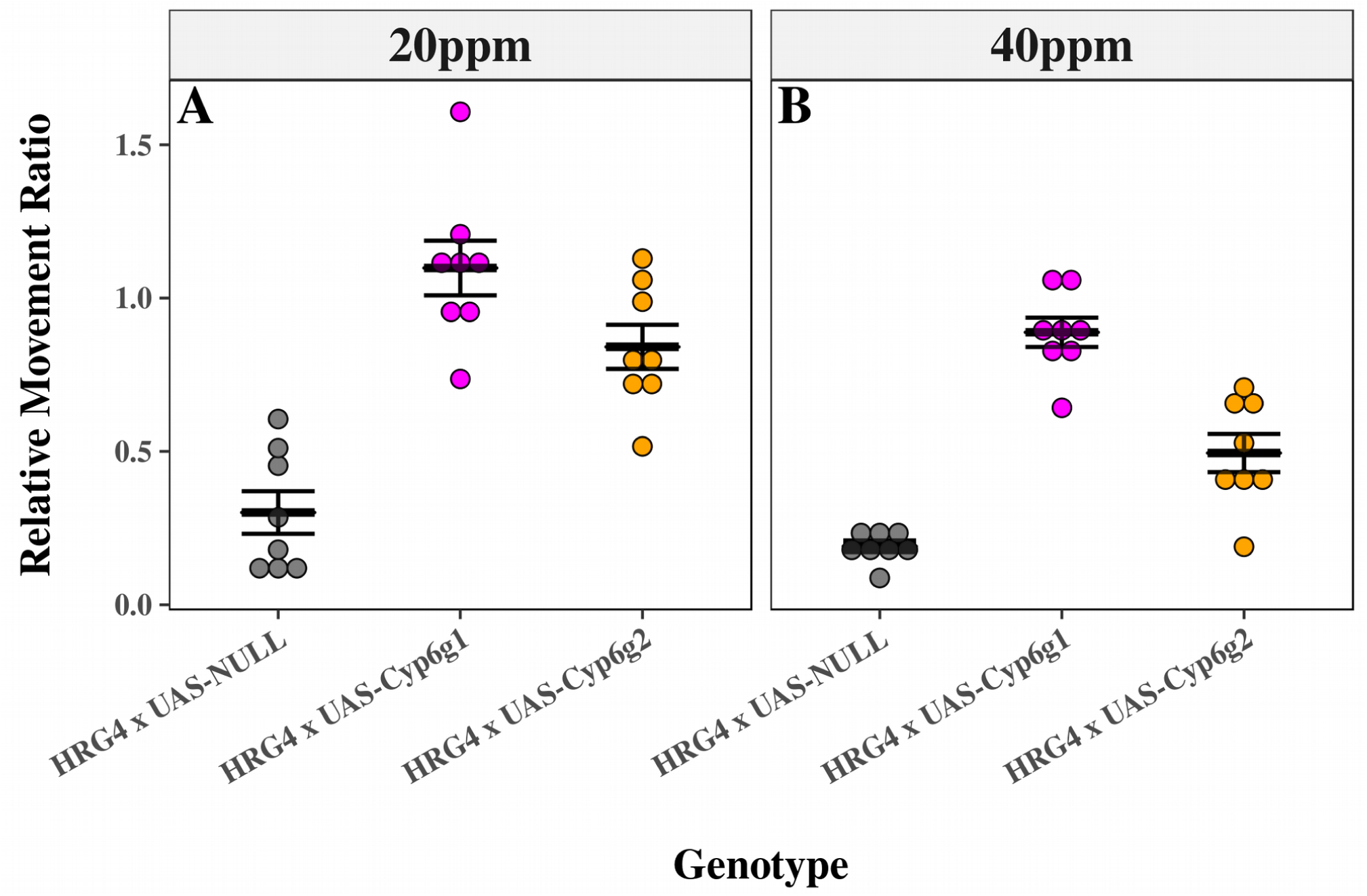
The Overexpression of Cyp6g1 and Cyp6g2 Confers Imidacloprid Resistance. The HR_GAL4 driver was used to overexpress Cyp6g1 (magenta) and Cyp6g2 (orange) from a common insertion site and their imidacloprid resistance was compared to their background control (grey). The Wiggle Index bioassay measured imidacloprid resistance and was performed at A) 20ppm and B) 40ppm in order to assess the magnitude of each gene’s ability to confer resistance. Error bars reflect the standard error of the mean. Significant differences were detected between all genotypes at all doses (ANOVA Tukey’s honestly significant difference) with p-values reported in Supplementary Table S6.

## Discussion

Of the candidates that were significantly (p<10^−5^) associated with imidacloprid response in the GWAS, several were in or near genes that have annotated roles in CNS development or function (Supplementary Table S3). Representative of this group is *Sickie,* the only candidate gene to be associated below the Bonferroni threshold (p<2.65x10^-8^). S*ickie* is orthologous to mammalian NAV2, which has been shown to regulate neuronal development^33^. Although originally identified as a regulator of *Relish* and posited to have a role in innate immune response^*34*^, *Sickie* has also been implicated in mushroom body development in *D. melanogaster*. Its expression is also highly enriched in the third instar larvae CNS, and genotypes carrying hypomorphic *Sickie* alleles showed axon growth defects^35^.

The mechanisms by which *Sickie* and other neuronal GWAS candidates might influence imidacloprid response are unknown. It may be that variation in such genes changes the amount of nAChRs at the synaptic membrane that can be targeted by imidacloprid. Alternatively, changing the connectivity of neurons could influence how imidacloprid’s signal is propagated after the insecticide has bound its target. Transgenic techniques are often used to explore the biological function of genes implicated in GWAS, but this was not performed in the current study. Many of these genes are homozygous lethal if knocked out. Furthermore, it is not clear that gene knockout, knockdown or overexpression would reproduce a resistance phenotype because these alleles would differ from those observed in the DGRP and may carry severe fitness costs. If the specific SNP associated with resistance was introduced via CRISPR, the difference may be too small to detect. However, the enrichment for neuronal genes in the GWAS candidate list suggests a role for CNS function in imidacloprid response. This may be a mechanism common to neurotoxic insecticides as a GWAS for azinphos-methyl resistance in the DGRP also implicated a high proportion of candidates enriched in the CNS (37.1% compared to 19.6% genome wide)^17^. The higher neuronal proportion of CNS enriched candidates (52.6%) in the current study may be due to the WI measuring a behavioural phenotype, that may be more strongly influenced by CNS function. It should be noted that variation in genes encoding the nAChR subunits known to be targeted by imidacloprid, *Dα1* and *Dβ2*^*5*^, was not shown to impact the insecticide response. This may be due to an absence of suitable variation in the DGRP. Studies with laboratory mutants suggest that resistance via these genes is associated with a loss of function that may involve a fitness cost^6^.

The capacity to metabolize imidacloprid also contributes to resistance in the DGRP; consideration of a subset of 9 of the most resistant and susceptible DGRP genotypes revealed that more imidacloprid metabolism occurred in resistant genotypes compared to susceptible ones (Fig. 2). However, such differences were only apparent in the media, unlike the RAL_517-Cyp6g1KO, which showed significantly different metabolite levels in the media and body at both one and six hour time points (Fusetto et. al this issue). This suggests that excretion is playing a critical role in the imidacloprid response within the DGRP. There appears to be a genetic component to excretion as the ratio between body and media metabolite levels varied between genotypes. In particular, RAL_509, the most susceptible genotype in this study, displays an interesting set of phenotypes. This genotype has a BA haplotype at the *Cyp6g1* locus, and produces high levels of both IMI-Olefin and IMI-5-OH levels, but these metabolites are disproportionately retained in the body (Fig. 2). Although the genetic basis for excretion was not explored in the current work, transporter proteins, such as ATP binding cassette transporters, have been implicated in insecticide resistance recently and could underpin differences in excretion between DGRP genotypes^36,37^. Further manipulation of transporters could reveal insights into how insecticides are excreted from the body.

The role played by metabolism in imidacloprid resistance was reinforced by the implication of *Cyp6g1* and *Cyp6g2* in the TWAS (Fig. 3; Supplementary Table S4). Although the involvement of *Cyp6g1* in imidacloprid resistance was known previously^13^, the magnitude of the contribution of *Cyp6g1* had not been tested. Removal of *Cyp6g1* from M haplotypes did not affect WI response, while knockouts in a BA haplotype reduced WI response and imidacloprid metabolism (Fig. 4, Fusetto et. al this issue). It may be that in M haplotypes there is not sufficient *Cyp6g1* to contribute to imidacloprid resistance. Previous studies have used RNAi to knock down the expression of *Cyp6g1* in wild-type backgrounds and seen either a slight increase in susceptibility to DDT or no change^38,39^. While lacking a clear understanding about the correlation between imidacloprid response and the amount of *Cyp6g1* expression, our data show that *Cyp6g1* does not contribute significantly to imidacloprid response in M haplotypes as measured with the WI. This does not rule out the possibility that such differences may be detected with other toxicological bioassays or with other insecticides.

The role of P450s apart from *Cyp6g1* in imidacloprid resistance was unknown in this species prior this study. Detection of the same metabolites produced by CYP6G1 (IMI-5-OH and IMI-Olefin) in RAL_517-Cyp6g1KO suggested that other P450s metabolize imidacloprid in that background (Fusetto et. al this issue). The ability of *Cyp6g2* to both metabolize and confer resistance to imidacloprid when transgenically expressed suggests that this gene may be one source of the residual resistance in RAL_517-Cyp6g1KO (Fig. 6; Supplementary Fig. S5). Based on evidence from the literature it is widely believed that there are two groups of P450s, those involved in metabolism and those involved in development. The *Cyp6g2* gene falls into both of these groups. In laboratory strains *Cyp6g2* is specifically expressed in the corpus allatum within the ring gland^40^ and is implicated in the synthesis of juvenile hormone^41^. However, ectopic expression of this gene in digestive tissues showed that it was able to confer resistance to nitenpyram and diazinon^32^. The current work extended the substrate specificity of CYP6G2 to imidacloprid, finding that it was able to produce the same metabolites as CYP6G1 and confer resistance (Fig. 6; Supplementary Fig. S5).

Although *Cyp6g2* expression was significantly associated with imidacloprid resistance in the TWAS, it was not known whether upregulation of this gene in resistant genotypes is restricted to its native ring gland specific expression pattern. Significantly higher levels of *Cyp6g2* expression in the digestive tissues of AA genotypes (Fig. 5E) suggests that this gene may contribute meaningfully to imidacloprid metabolism within the DGRP. This increase is at least partially tissue specific, as increases in the digestive tissues were far higher than those in whole adult bodies (Fig. 5B,E). The most parsimonious source of the change in expression level and pattern is the presence of an *Accord* element, which could act as an enhancer, increasing the expression of both *Cyp6g1* and *Cyp6g2* in the digestive tissues. *Cyp6g2* may represent the limit of *Accord’s* range as expression of the more distant *Cyp6t3* did not appear to be influenced by the presence of *Accord*. The regulation of *Cyp6g2* by *Accord* is not the only possible mechanism for of the observed expression in metabolic tissue. Other differences between AA and M haplotypes could influence *Cyp6g2* expression and the expression pattern in BA haplotypes is unknown. Small sample sizes and the consideration of only AA and M haplotypes preclude the establishment of any definitive mechanism for regulating *Cyp6g2* expression.

Much still remains unresolved about the relative contribution of different alleles to complex phenotypes such as insecticide resistance. While far more attention has been given to cases of monogenic resistance, all populations reflect a distribution of resistance levels among individuals that is governed by many loci^2^. This is true even in populations where resistance alleles have gone to fixation. Variation at the *Cyp6g1* locus contributes significantly to imidacloprid resistance in the DGRP and is likely the largest single factor in determining the likelihood an insect survives an exposure. However, while removal of *Cyp6g1* from the resistant RAL_517 genotype increased susceptibility to imidacloprid, the RAL_517-Cyp6g1KO genotype was still more resistant than approximately half of the DGRP genotypes (Fig. 4A). This indicates that *Cyp6g1* significantly contributes to imidacloprid resistance, but also highlights the supporting role played by other genes such as *Cyp6g2* and *Sickie* which likely contribute in smaller ways. Imidacloprid resistance in the DGRP can then be thought of as polygenic but with a single gene making a contribution far larger than the rest.

## Methods

### Fly Genotypes

All genotypes used in this study were ordered from the Bloomington Drosophila Stock Center (Bloomington, Indiana) or generated in this study (Supplementary Table S7). 178 of the 201 DGRP genotypes was tested in the the initial WI screen. A subset of 9 of the most imidacloprid resistant and susceptible DGRP genotypes were chosen for imidacloprid metabolism analysis via HPLC-MS. The Actin-Cas9 genotype (Bloomington #54590) and a genotype containing an attP landing site (attP40; 2L:5,108,448:5,108,448; Bloomington #25709) were used for used to knock out the *Cyp6g1* gene in three different genetic backgrounds. Canton-S-Cyp6g1KO was generated in the Canton-S background, Wxac-Cyp61KO was generated from the Wxac background (Actin-Cas9 with the X chromosome replaced with the one from the w^1118^ genotype). RAL_517-Cyp6g1KO was created in the RAL_517 background, a genotype chosen due to its high level of imidacloprid resistance and *Cyp6g1* expression among DGRP genotypes. RAL_517 carries a BA haplotype while Canton-S and Wxac both carry M haplotypes and were far more susceptible to imidacloprid.

The three *Cyp6g1* knockouts generated here were created by using a transgenic CRISPR strategy described recently^42^. Briefly, Cyp6g1-sgRNA plasmids were made by first cutting the PCFD4 plasmid (Addgene #49411) with the restriction enzyme Bsb1. A separate fragment was generated by amplifying a portion of the PCFD4 plasmid with the Cyp6g1-PCFD4 primer set (Supplementary Table S8; 60°C annealing, 1 minute extension), introducing *Cyp6g1* sgRNAs into the PCR product. The cut plasmid and PCR product were then reassembled using the Gibson assembly kit (New England Biolabs) to make a circular PCFD4 plasmid with two *Cyp6g1* sgRNAs under the control of two U6 promoters. Verification of this modification was accomplished by sequencing the plasmid using the PCFD4_seq primer (Supplementary Table S8).

This plasmid was then injected into a genotype expressing φ-31 integrase and which contained an attP landing site (attP40; 2L:5,108,448:5,108,448; Bloomington #25709), both of which facilitated the integration of the modified PCFD4 into the germline. Transgenic flies were identified by scoring the visible marker vermilion eyes which was restored to wild type upon successful PCFD4 integration. Chromosomes from Actin-Cas9 and the sgRNA expressing genotype were brought together in a crossing scheme which made near identical deletions of *Cyp6g1* in each background (Supplementary Fig. S6A). The resulting deletion was confirmed by amplifying across the *Cyp6g1* deletion using the Cyp6g1-KO primer set (Supplementary Table S8; 56°C, 2 minutes extension). Sequencing this PCR product revealed the almost complete removal of the gene (Supplementary Fig. S6B), and the remaining transcript was predicted at 73 amino acids long (reduced from 524). Failure to amplify any full copies of the gene indicated that both copies of *Cyp6g1* present in this RAL_517 were successfully deleted.

Overexpression of the genes *Cyp6g1* and *Cyp6g2* was achieved in the midgut Malpighian tubules and the fat body using the GAL4/UAS system^43^ and the HR_GAL4 driver^14^. In testing the capacity of CYP6G2 to metabolize imidacloprid, a previously reported UAS-Cyp6g2-3d genotype was used^32^. However, this genotype was created using a random insertion method, precluding a direct comparison of the impact of *Cyp6g1* and *Cyp6g2* on resistance. Hence, new UAS-Cyp6g1 and UAS-Cyp6g2 genotypes that share a common insertion site (attP40) were generated in this study. Open reading frames from each gene were amplified from the *w*^*1118*^ background using Q5 high fidelity master mix (New England Biolabs). Each PCR fragment was A-tailed and cloned into the PGEM T-easy vector (Promega). The fragments were then cut out of this vector using the NotI enzyme and ligated into the pUASt-attB vector^44^. Plasmids were injected into a genotype carrying the same attP landing site genotype used for knockouts (attP40) and an X chromosome with a white^-^allele (Bloomington stock # 24749). These two genotypes were compared to their control, which was created by injecting an empty pUASt-attB vector into the same background.

### The Wiggle Index

The response of *D. melanogaster* larvae to imidacloprid was measured using the WI assay, which estimates the insecticidal effect by quantifying the reduction in motility during insecticide exposure^31^. Third instar larvae of each genotype were picked, 25 per well, into a NUNC cell culture treated 24 well plate (Thermo-Scientific) preloaded with 200µL 5% w/v sucrose (AR Grade; Chem Supply) in distilled H_2_0. Larvae were filmed for 30 seconds at two time points, 0min (before starting the exposure) and at 60min after the addition of specific concentrations of imidacloprid (200 g L^-1^ Confidor®; Bayer Crop Science). Subsequently, the WI ImageJ script was used to quantify the total motion in each well at each time point. The ratio between initial and final motility was used to calculate Relative Movement Ratios (RMRs), which were averaged to estimate imidacloprid response for each genotype tested in this study. 178 DGRP genotypes were considered at doses of 25 and 100ppm as was RAL_517 and RAL_517-Cyp6g1KO. Other *Cyp6g1* knockouts were tested at 5ppm. UAS-Cyp6g1 and UAS-Cyp6g2 were tested at 20 and 40ppm.

### Analysis of the WI and Candidate Gene Selection

Broad sense heritability (H^2^) of imidacloprid resistance was measured by comparing the variance of RMRs within genotypes to the variance between genotypes. The association of initial motility values (at 0 minutes) with final RMR was also tested in order to test the any confounding effects of starting movement on imidacloprid response. The correlation between 25 and 100ppm RMRs was tested to observe the relationship between the two phenotypes. All these associations were tested by assessing the fit to a linear model using Pearson’s correlation test.

In order to implicate individual QTLs in imidacloprid resistance, mean RMRs for each genotype were used as input data for the Mackay lab DGRP GWAS pipeline^28^ found at (http://dgrp2.gnets.ncsu.edu/). Responses to the two doses (25 and 100ppm) were considered as separate phenotypes. Similarly, a transcriptome wide association study, was performed by associating the phenotypic Wiggle data with expression data recently generated in 185 DGRP genotypes^30^. Analysis was performed using a modification of a recently reported pipeline^17^, which tested the fit of linear models (Pearson’s correlation test) correlating the expression level of an individual transcript and mean RMR among genotypes. Transcript levels for each gene were averaged between males and females as larvae were not sexed before testing in the WI assay. Furthermore, the expression quantitative trait loci (eQTLs) of each significant transcript^30^ were cross examined with significant SNPs from the GWAS in order to test if significantly associated genomic loci regulate the expression of any significantly associated transcripts.

Candidate genes from the GWAS and TWAS were chosen based on their significance of association with either the 25 or the 100ppm phenotype. Any GWAS variant eliciting a p-value of less than 10^−-5^ and any transcript with a p-value of less than 10^−3^ was considered as a candidate in the current study. In the case of the GWAS, variants were assigned to their nearest annotated gene according to the software SnpEff (built into the Mackay pipeline), and only p-values derived from simple regression were considered (mixed models were not considered). Candidate genes were further examined by testing enrichment in third instar larval tissues using the modEncode transcriptome datasets^45^.

### Quantification of Imidacloprid and its Metabolites

Levels of imidacloprid and its metabolites were quantified via HPLC-MS following the method described by Fusetto et. al (this issue). Quadruplicates of 200 third instar larvae from a given genotype were placed into 200µL of 5% analytical reagent sucrose and were exposed to a 50:50 mix of ^12^C_6_:^13^C_6_ imidacloprid (99% analytical reagent) at a concentration of 25ppm for 1 hour. Larvae were then recovered from the solution and washed 3 times with 3mL of distilled H_2_O to remove chemical from the cuticle, and the exposure media was collected separately in order to estimate the amount of each chemical excreted. The compounds in each sample were quantified using high-performance liquid chromatography (HPLC) coupled with an Agilent 6520 Q-TOF Mass Spectrometer (Agilent Technologies, Inc., Santa Clara, CA, USA). These measurements were taken for a subset of the 9 most resistant and susceptible DGRP lines and the UAS-Cyp6g2-3d genotype^32^ as well as all relevant control lines. In the case of the DGRP subset, the correlation between the mean of each metabolic phenotype (each chemical in larval bodies or excreta) and the mean of each genotype’s WI RMR at 25ppm (Fig. 1A) was tested among genotypes using Pearson’s correlation test. All other comparisons were made using a students t-test to compare genotypes, with a Bonferroni correction applied to correct for multiple testing. Values was reported in either parts per billion (ppb) or, in the case of UAS-Cyp6g2-3d, the area under the chromatogram peak was used as a relative quantity of the chemical.

### Measurement of Cyp6g1 and Cyp6g2 Expression in Digestive Tissues

The expression of *Cyp6g1, Cyp6g2* and *Cyp6t3* was measured in digestive tissues (midgut and Malpighian tubules) of a subset of DGRP lines. 4 lines were chosen in total. Two carried the AA haplotype (RAL_138, RAL_360) and two carried the M (lines RAL_776, RAL_843). 20 midguts and the associated malpighian tubules from each genotype were used for a single biological replicate and 5 replicates were taken from each genotype. RNA was extracted from each sample using the TriSure (Bioline) protocol, and cDNA was generated using the Superscript III reverse transcriptase (NEB). All primers and qPCR parameters were reported previously^32^. Expression was quantified using the 2^−ΔΔ CT^ method and normalized to the level of the genotype showing the lowest expression level tested (RAL_776). Differences in expression among genotypes were compared using analysis of variance (ANOVA) with Tukey’s honestly significant difference post-hoc test.

### Data Analysis and Availability

All raw data generated in this study is available for download at https://github.com/shanedenecke/Denecke_et_al_2017_Imidacloprid_DGRP and was analysed in R^46^. An accompanying R markdown document, also available on Rpubs (http://rpubs.com/shanedenecke/Denecke_DGRP). This document provides R code to obtain the raw data and reproduce all the figures and tables presented in this study. Supplementary Fig. S7 and S8 are not included in the markdown document because these do not contain analysis driven information. Unless otherwise stated all p values represent Student’s t-tests (for pairwise comparisons), ANOVA with Tukey’s post-hoc test (3 comparisons of three or more variables) and Pearson’s correlation test (for correlations and linear regressions).

## Acknowledgements

The authors would like to thank the University of Melbourne for the *Writing Up Scholarship* and the Melbourne International Research Scholarship that assisted in financing this work. Furthermore, we must acknowledge Hang Nuoc Bao Hang for assisting in larvae picking. Llewellyn Green and Trent Perry also contributed productive discussions on the topic matter. Lastly, we must thank Michael Murray of the University of Melbourne for providing fly facilities during the renovations of our own flyroom.

## Author Contributions

S.D and Ph.B: Writing the manuscript

S.D, Ph.B, R.F, Pa.B, A.F-L: Editing the manuscript

S.D, R.F, F.M, A.G: Laboratory Work

S.D, R.F, A.F-L, Pa.B: Analysis of data

Ph.B, R.O’H: Funding and overall project design

---

**Figure S1-.**
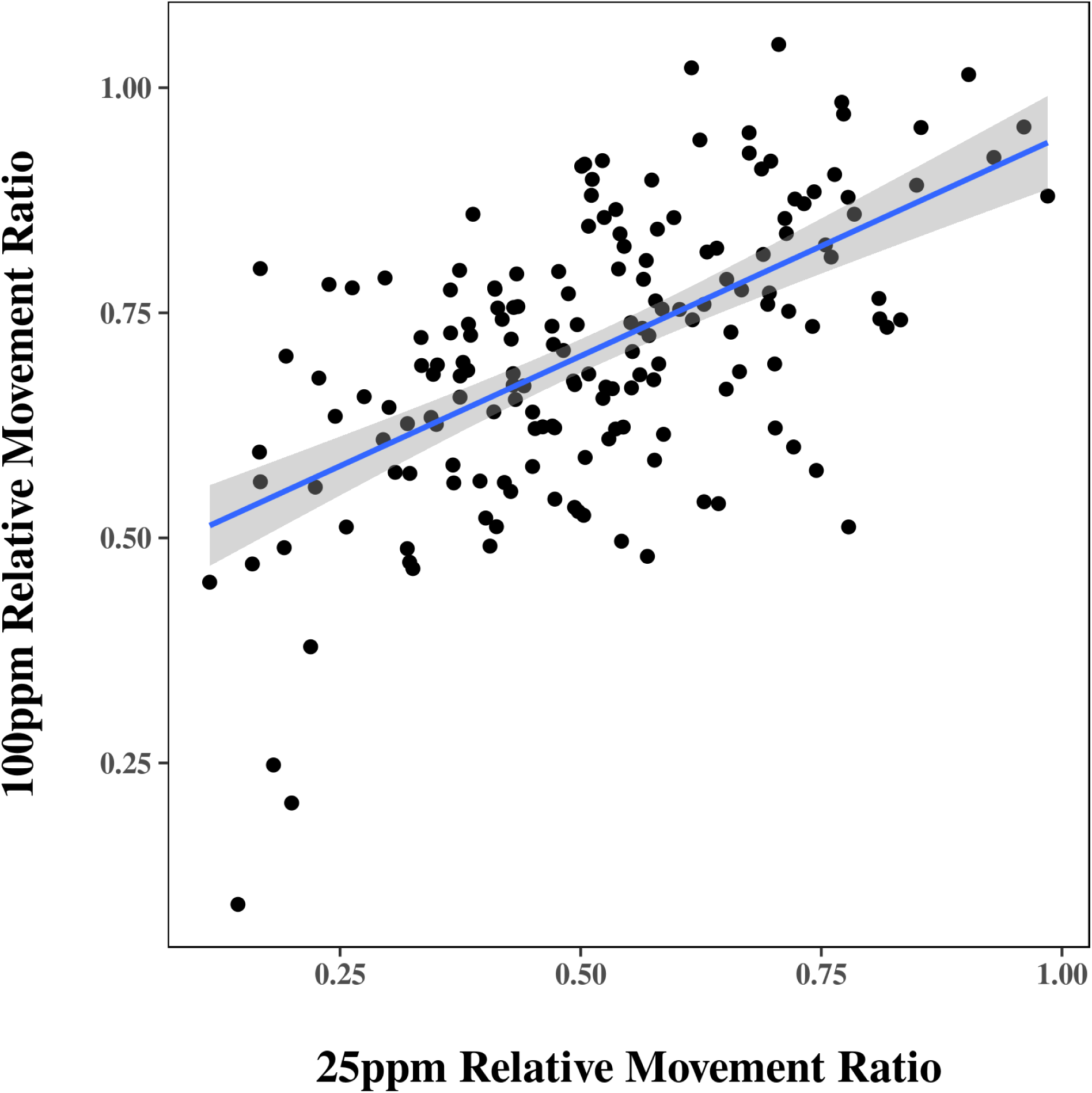
The Correlation of 25 and 100ppm Imidacloprid Response. For each genotype the 25 and 100ppm mean RMR values were plotted and a linear regression line was fit, showing substantial correlation. The shaded area reflects the 95% confidence interval of the linear model.

**Figure S2-.**
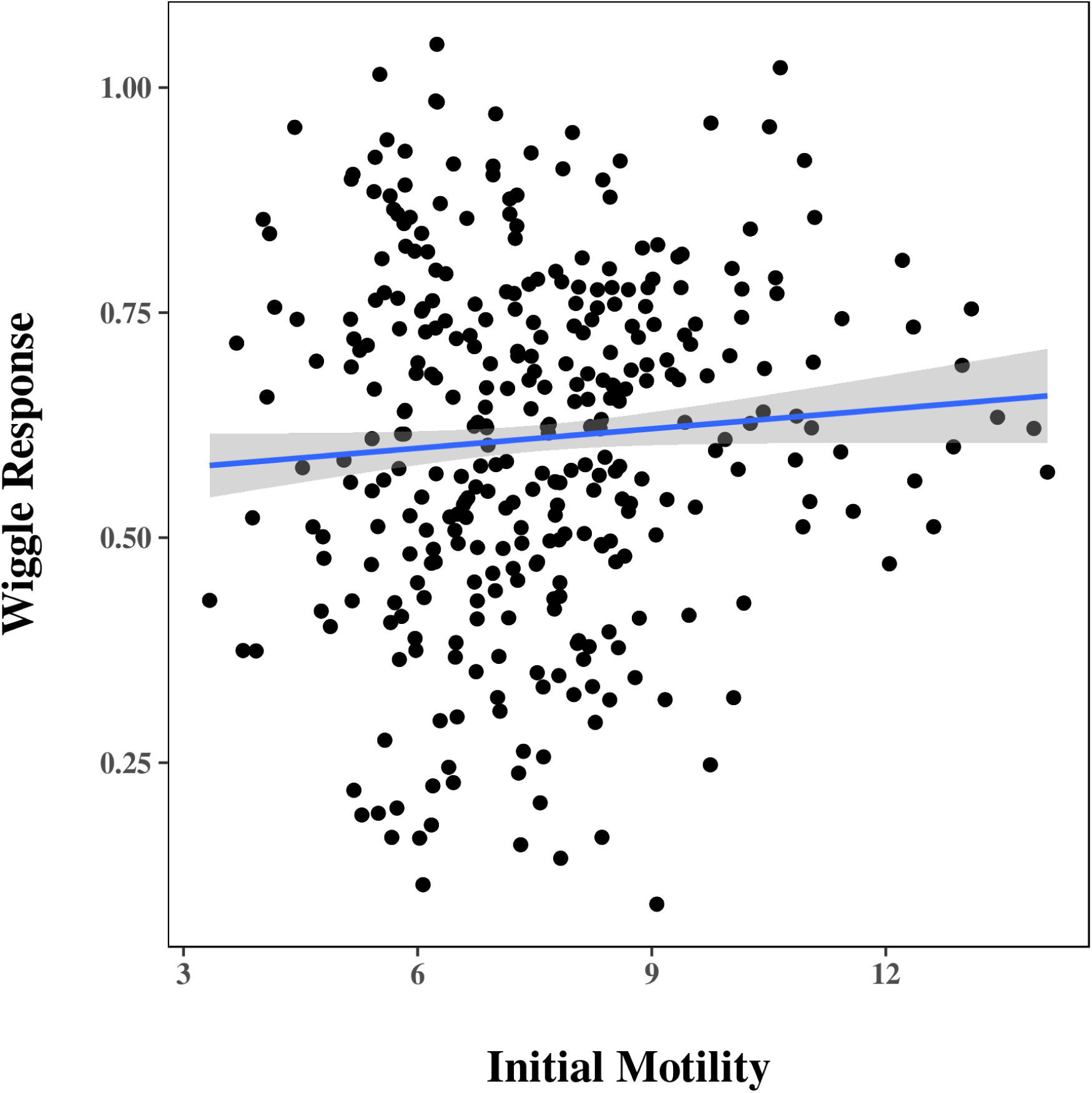
The Correlation of Initial Motility with Imidacloprid Response. Raw Wiggle Index values at time 0 (x axis) were compared to RMR values after 60 minutes (y axis), suggesting no correlation. The shaded area reflects the 95% confidence interval for the linear model.

**Figure S3-.**
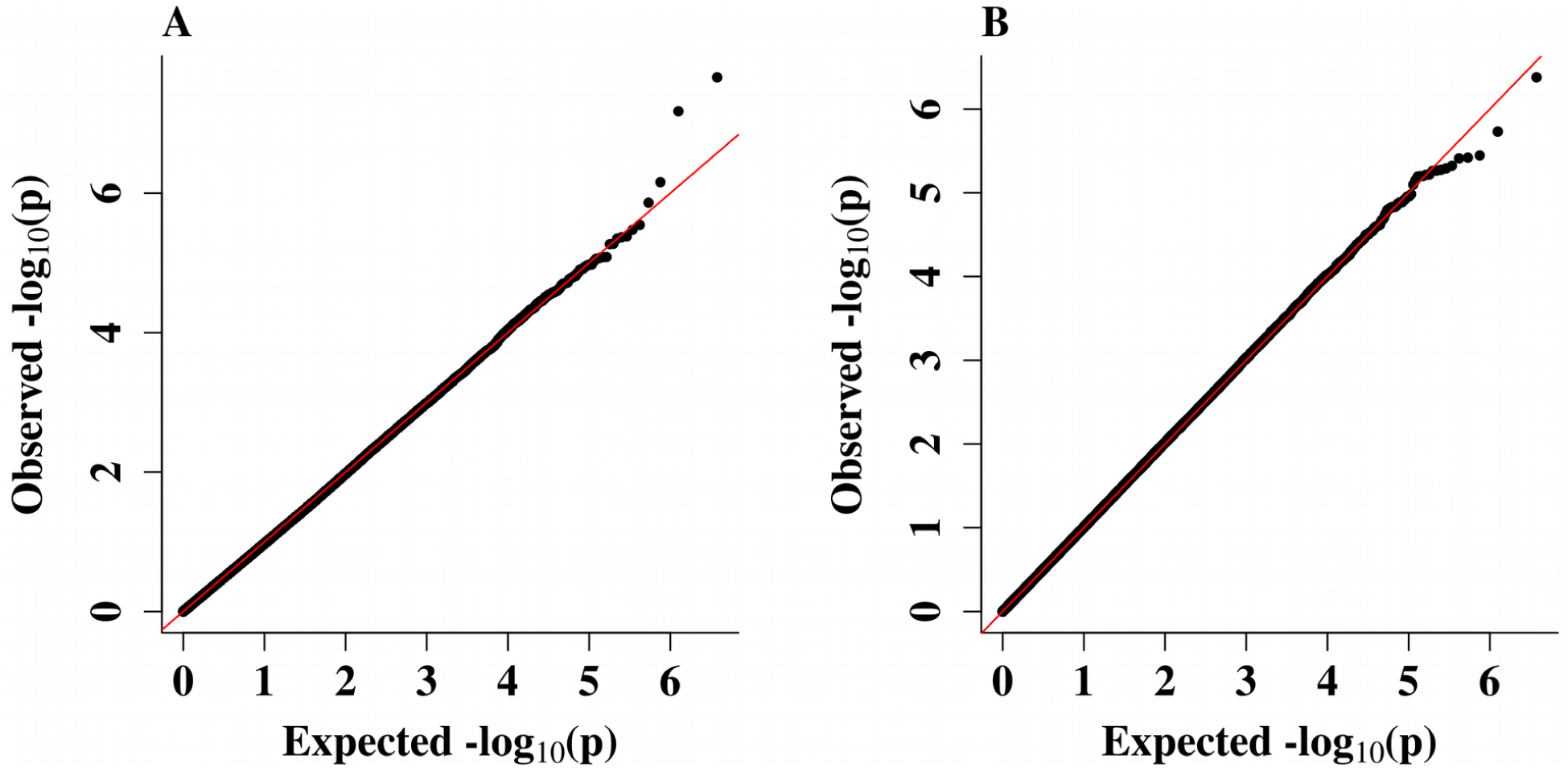
QQ Plots. Quantile Quantile (QQ) plots were made for the GWAS using A) 25ppm and B) 100ppm RMRs. Each plot shows the expected versus the observed distribution of the significance (p-values) of each annotated variant. The red line indicates where the observed values should fall if they were to match the expected values exactly.

**Figure S4-.**
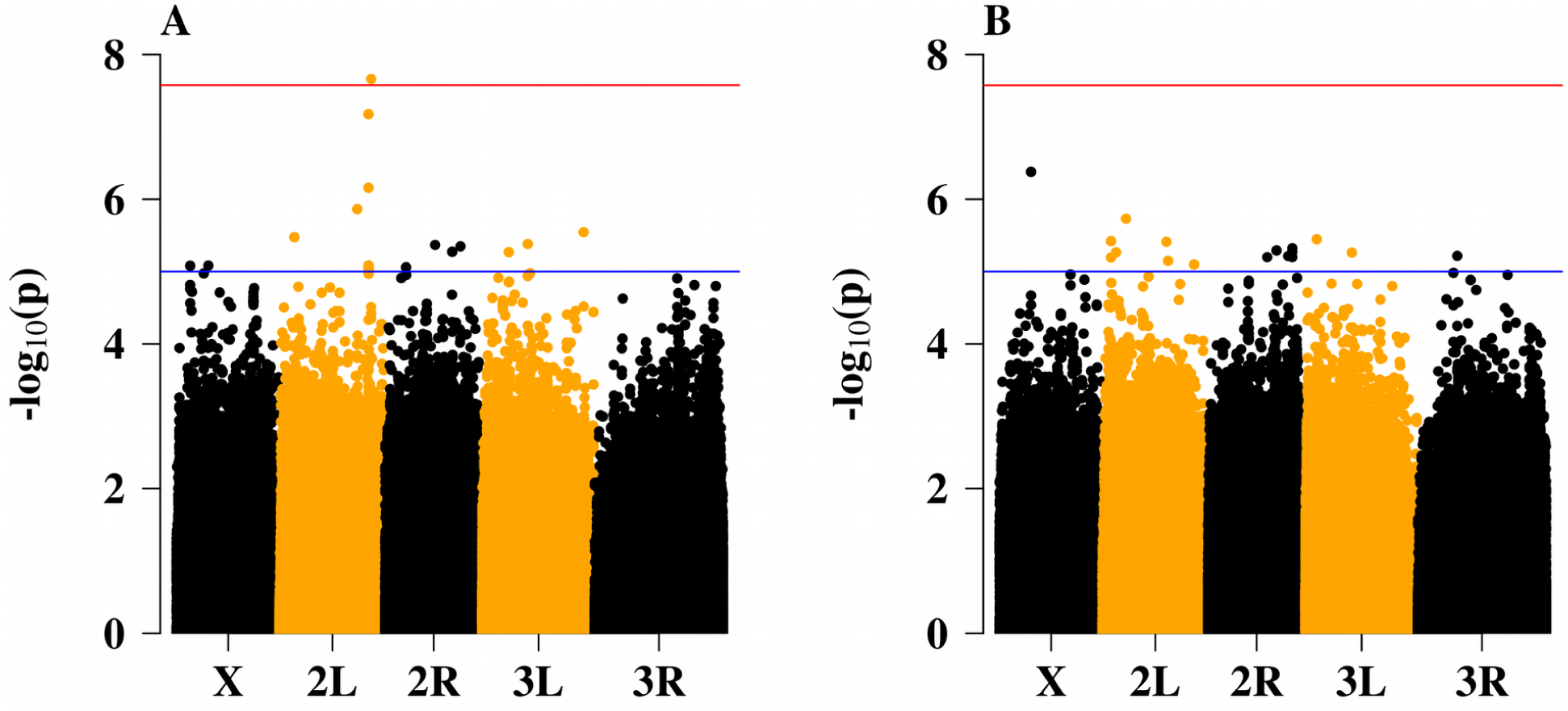
Manhattan Plots. Manhattan plots were made for the GWAS using A) 25ppm and B) 100ppm RMR values. Each plot shows the significance of a p-value on the Y axis and the position of the genetic variant on the X axis. Genome wide significance thresholds are shown at p=10^−5^ and p=2.65x10^−7^ (Bonferroni threshold)

**Figure S5-.**
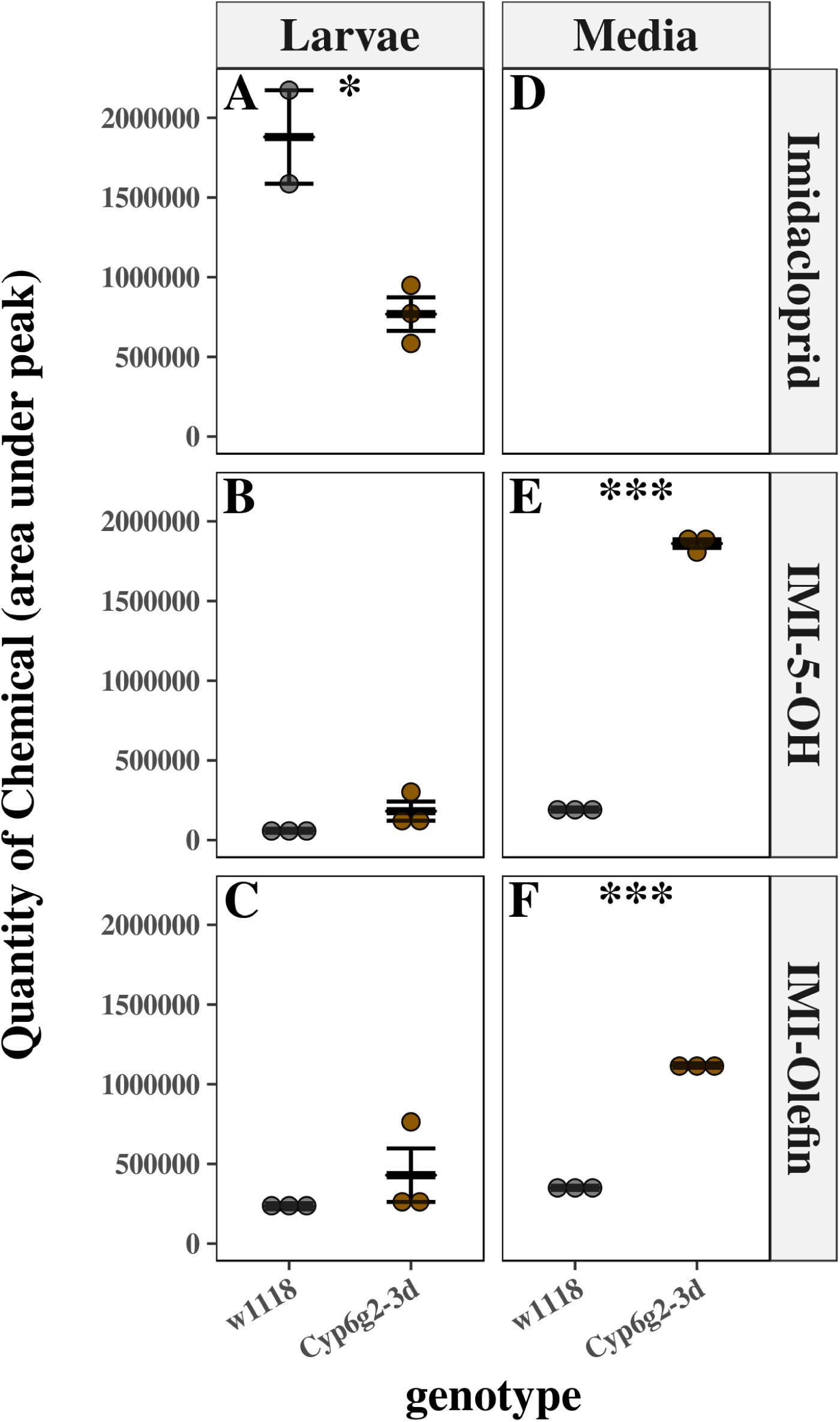
Imidacloprid Metabolism in UAS-Cyp6g2-3d. The amount of Imidacloprid (A,D) IMI-5-OH (B,E) and IMI-Olefin (C,F) recovered from larval bodies (A-C) or the media (D-F) is reported in HR-GAL4 x w^1118^ (grey) and HR-GAL4 x UAS-Cyp6g2 (brown). No data is presented for imidacloprid in the media, due to the relative abundance of this molecule in the media, which makes detecting changes impossible. Error bars represent standard error of the mean. Stars represent the significance of the difference between the two genotypes measured by he Students T-test corrected for multiple comparisons with the Bonferroni method (*=p<.05; **=p<.001; ***=p<.0001).

**Figure S6-.**
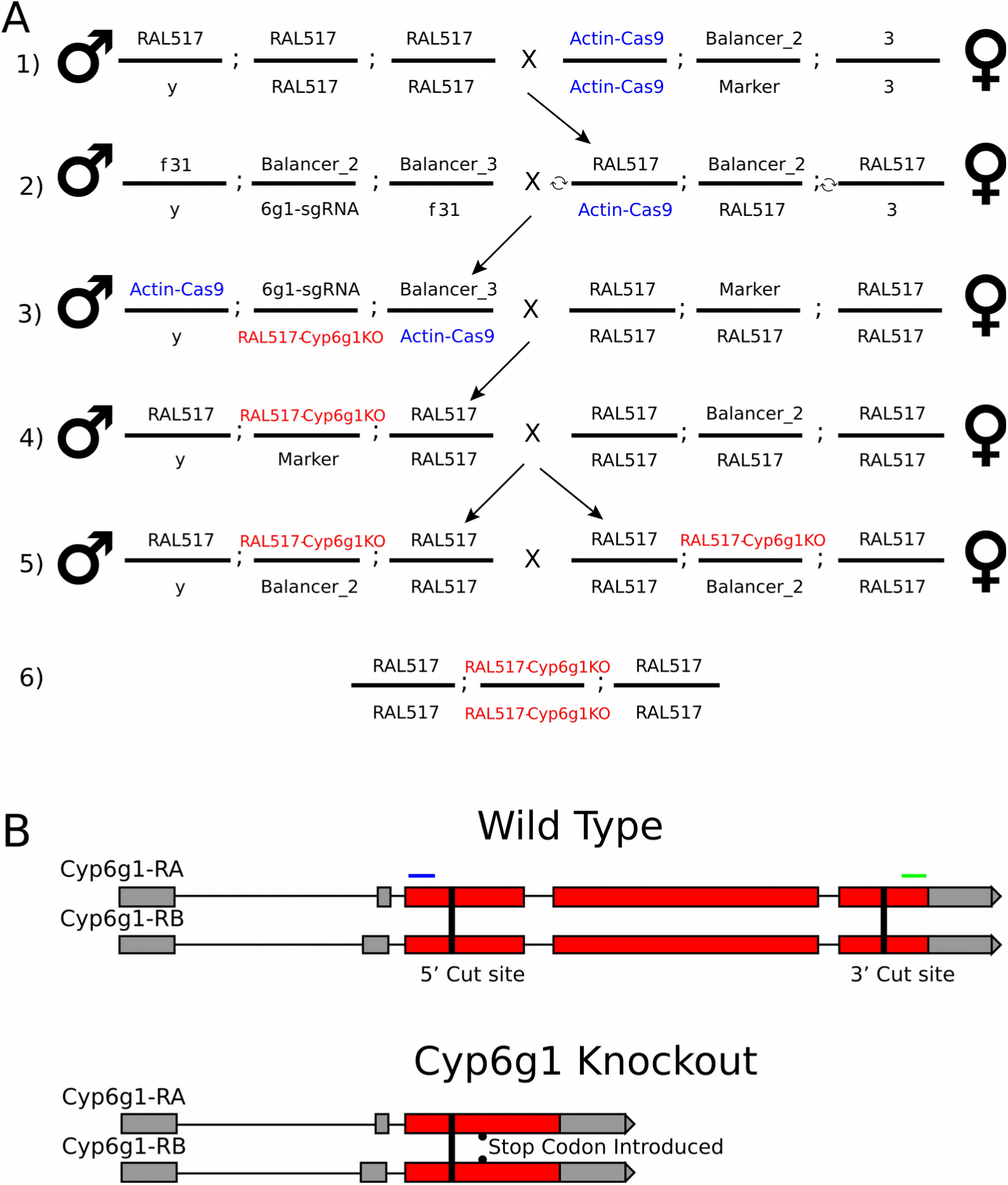
Crossing scheme and Cyp6g1 knock-out. (A) In step 1, the RAL517 line was crossed to a line that expresses Cas9 under the control of the Actin promoter (Actin-Cas9) with a balanced 2nd chromosome. The Actin-Cas9 cassette is inserted on the X chromosome (highlighted in blue). In step 2, males from the line carrying the balanced Cyp6g1-sgRNA cassette were crossed to females from cross 1. In step 3, backcross males from cross 2 were crossed into a balanced RAL517 background. Because Cas9 and Cyp6g1-sgRNAs are simultaneously expressed in males from cross 2, the deletion of the Cyp6g1 gene will occur at an appreciable frequency. In step 4, males from cross 3 were backcrossed again into a balanced copy of the RAL517 background. This made the X chromosome homozygous for RAL517 and put the deletion (RAL517-Cyp6g1KO) over the CyO balancer. In step 5, males and females from cross 4 carrying RAL517 Cyp6g1 2nd chromosome over the CyO balancer chromosome were crossed together. 6) The homozygous RAL517-Cyp6g1KO flies were identified as the progeny of the cross 5 that appear wild type. indicates recombination. (B) Schematic representation of the Cyp6g1 knock-out outlines that both copies of Cyp6g1 gene are removed from the RAL517 genome. Red boxes indicates coding sequences. The position of the Cyp6g1KO_F and Cyp6g1KO_R primers are reported in blue and green respectively

## Bibliography

1. ffrench-Constant, R. H. The Molecular Genetics of Insecticide Resistance. Genetics 194, (2013).

2. McKenzie, J. A. & Batterham, P. The genetic, molecular and phenotypic consequences of selection for insecticide resistance. Trends Ecol. Evol. 9, 166–9 (1994).

3. Groeters, F. R. & Tabashnik, B. E. Roles of Selection Intensity, Major Genes, and Minor Genes in Evolution of Insecticide Resistance. J. Econ. Entomol. 93, 1580–1587 (2000).

4. Bai, D., Lummis, S. C. R., Leicht, W., Breer, H. & Sattelle, D. B. Actions of imidacloprid and a related nitromethylene on cholinergic receptors of an identified insect motor neurone. Pestic. Sci. 33, 197–204 (1991).

5. Perry, T., Heckel, D. G., McKenzie, J. A. & Batterham, P. Mutations in Dα1 or Dβ2 nicotinic acetylcholine receptor subunits can confer resistance to neonicotinoids in *Drosophila melanogaster*. Insect Biochem. Mol. Biol. 38, 520–8 (2008).

6. Somers, J., Luong, H. N. B., Mitchell, J., Batterham, P. & Perry, T. Pleiotropic Effects of Loss of the Dα1 Subunit in *Drosophila melanogaster*: Implications for Insecticide Resistance. Genetics 205, 263–271 (2017).

7. Bass, C., Denholm, I., Williamson, M. S. & Nauen, R. The global status of insect resistance to neonicotinoid insecticides. Pestic. Biochem. Physiol. 121, 78–87 (2015).

8. Feyereisen, R. Insect P450 enzymes. Annu. Rev. Entomol. 44, 507–33 (1999).

9. Karunker, I. et al. Over-expression of cytochrome P450 CYP6CM1 is associated with high resistance to imidacloprid in the B and Q biotypes of *Bemisia tabaci* (Hemiptera: Aleyrodidae). Insect Biochem. Mol. Biol. 38, 634–644 (2008).

10. Puinean, A. M. et al. Amplification of a cytochrome P450 gene is associated with resistance to neonicotinoid insecticides in the aphid *Myzus persicae*. PLoS Genet. 6, e1000999 (2010).

11. Bass, C. et al. Overexpression of a cytochrome P450 monooxygenase, CYP6ER1, is associated with resistance to imidacloprid in the brown planthopper, *Nilaparvata lugens*. Insect Mol. Biol. 20, 763–73 (2011).

12. Le Goff, G. & Hilliou, F. Resistance evolution in Drosophila: the case of CYP6G1. Pest Manag. Sci. (2016). doi:10.1002/ps.4470

13. Daborn, P. J. A Single P450 Allele Associated with Insecticide Resistance in Drosophila. Science (80-.). 297, 2253–2256 (2002).

14. Chung, H. et al. Cis-regulatory elements in the Accord retrotransposon result in tissue-specific expression of the *Drosophila melanogaster* insecticide resistance gene Cyp6g1. Genetics 175, 1071–7 (2007).

15. Catania, F. et al. World-wide survey of an Accord insertion and its association with DDT resistance in *Drosophila melanogaster*. Mol. Ecol. 13, 2491–504 (2004).

16. Schmidt, J. M. et al. Copy number variation and transposable elements feature in recent, ongoing adaptation at the *Cyp6g1* locus. PLoS Genet. 6, e1000998 (2010).

17. Battlay, P., Schmidt, J. M., Fournier-Level, A. & Robin, C. Genomic and Transcriptomic Associations Identify a New Insecticide Resistance Phenotype for the Selective Sweep at the *Cyp6g1* Locus of *Drosophila melanogaster*. G3; Genes|Genomes||Genetics 6, 2573–2581 (2016).

18. Ford, K. A. & Casida, J. E. Unique and common metabolites of thiamethoxam, clothianidin, and dinotefuran in mice. Chem. Res. Toxicol. 19, 1549–56 (2006).

19. Ford, K. A. & Casida, J. E. Comparative metabolism and pharmacokinetics of seven neonicotinoid insecticides in spinach. J. Agric. Food Chem. 56, 10168–75 (2008).

20. Pandey, G., Dorrian, S. J., Russell, R. J. & Oakeshott, J. G. Biotransformation of the neonicotinoid insecticides imidacloprid and thiamethoxam by *Pseudomonas sp. 1G*. Biochem. Biophys. Res. Commun. 380, 710–4 (2009).

21. Joussen, N., Heckel, D. G., Haas, M., Schuphan, I. & Schmidt, B. Metabolism of imidacloprid and DDT by P450 CYP6G1 expressed in cell cultures of *Nicotiana tabacum* suggests detoxification of these insecticides in Cyp6g1-overexpressing strains of *Drosophila melanogaster*, leading to resistance. Pest Manag. Sci. 64, 65–73 (2008).

22. Hoi, K. K. et al. Dissecting the insect metabolic machinery using twin ion mass spectrometry: a single P450 enzyme metabolizing the insecticide imidacloprid *in vivo*. Anal. Chem. 86, 3525–32 (2014).

23. Riaz, M. A. et al. Molecular mechanisms associated with increased tolerance to the neonicotinoid insecticide imidacloprid in the dengue vector *Aedes aegypti*. Aquat. Toxicol. 126, 326–37 (2013).

24. Ilias, A. et al. Transcription analysis of neonicotinoid resistance in Mediterranean (MED) populations of *B. tabaci* reveal novel cytochrome P450s, but no nAChR mutations associated with the phenotype. BMC Genomics 16, 939 (2015).

25. King-Jones, K., Horner, M. A., Lam, G. & Thummel, C. S. The DHR96 nuclear receptor regulates xenobiotic responses in *Drosophila*. Cell Metab. 4, 37–48 (2006).

26. Misra, J. R., Horner, M. A., Lam, G. & Thummel, C. S. Transcriptional regulation of xenobiotic detoxification in *Drosophila*. Genes Dev. 25, 1796–806 (2011).

27. Mackay, T. F. C. et al. The *Drosophila melanogaster* Genetic Reference Panel. Nature 482, 173–8 (2012).

28. Huang, W. et al. Natural variation in genome architecture among 205 *Drosophila melanogaster* Genetic Reference Panel lines. Genome Res. 24, 1193–208 (2014).

29. Ayroles, J. F. et al. Systems genetics of complex traits in *Drosophila melanogaster*. Nat. Genet. 41, 299–307 (2009).

30. Huang, W. et al. Genetic basis of transcriptome diversity in Drosophila melanogaster. Proc. Natl. Acad. Sci. U. S. A. 112, E6010–9 (2015).

31. Denecke, S., Nowell, C. J., Fournier-Level, A., Perry, T. & Batterham, P. The wiggle index: An open source bioassay to assess sub-lethal insecticide response in *Drosophila melanogaster*. PLoS One 10, e0145051 (2015).

32. Daborn, P. J. et al. Evaluating the insecticide resistance potential of eight *Drosophila melanogaster* cytochrome P450 genes by transgenic over-expression. Insect Biochem. Mol. Biol. 37, 512–9 (2007).

33. Marzinke, M. A., Mavencamp, T., Duratinsky, J. & Clagett-Dame, M. 14-3-3ε and NAV2 interact to regulate neurite outgrowth and axon elongation. Arch. Biochem. Biophys. 540, 94–100 (2013).

34. Foley, E. et al. Functional Dissection of an Innate Immune Response by a Genome-Wide RNAi Screen. PLoS Biol. 2, e203 (2004).

35. Abe, T. et al. Correction: the NAV2 homolog *Sickie* regulates F-actin-mediated axonal growth in *Drosophila* mushroom body neurons via the non-canonical Rac-Cofilin pathway. Development 142, 1021 (2015).

36. Dermauw, W. & Van Leeuwen, T. The ABC gene family in arthropods: comparative genomics and role in insecticide transport and resistance. Insect Biochem. Mol. Biol. 45, 89–110 (2014).

37. Merzendorfer, H. ABC Transporters and Their Role in Protecting Insects from Pesticides and Their Metabolites. Advances in Insect Physiology 46, (Elsevier, 2014).

38. Yang, J. et al. A Drosophila systems approach to xenobiotic metabolism. Physiol. Genomics 30, 223–31 (2007).

39. McCart, C. & Ffrench-Constant, R. H. Dissecting the insecticide-resistance-associated cytochrome P450 gene Cyp6g1. Pest Manag. Sci. 64, 639–45 (2008).

40. Chung, H. et al. Characterization of *Drosophila melanogaster* cytochrome P450 genes. Proc. Natl. Acad. Sci. U. S. A. 106, 5731–6 (2009).

41. Christesen, D. et al. Transcriptome Analysis of *Drosophila melanogaster* Third Instar Larval Ring Glands Points to Novel Functions and Uncovers a Cytochrome p450 Required for Development. G3 (Bethesda). 7, 467–479 (2017).

42. Port, F., Chen, H.-M., Lee, T. & Bullock, S. L. Optimized CRISPR/Cas tools for efficient germline and somatic genome engineering in *Drosophila*. Proc. Natl. Acad. Sci. U. S. A. 111, E2967–76 (2014).

43. Brand, A. H. & Perrimon, N. Targeted gene expression as a means of altering cell fates and generating dominant phenotypes. Development 118, 401–15 (1993).

44. Bischof, J., Maeda, R. K., Hediger, M., Karch, F. & Basler, K. An optimized transgenesis system for Drosophila using germ-line-specific phiC31 integrases. Proc. Natl. Acad. Sci. U. S. A. 104, 3312–7 (2007).

45. ModEncode Consortium, et al. Identification of functional elements and regulatory circuits by Drosophila modENCODE. Science 330, 1787–97 (2010).

46. R Core Team. R: A Language and Environment for Statistical Computing. (2017)

